# A simple genetic architecture and low constraint allows rapid floral evolution in a diverse and recently radiating plant genus

**DOI:** 10.1101/516377

**Authors:** Jamie L. Kostyun, Matthew J.S. Gibson, Christian M. King, Leonie C. Moyle

## Abstract

- Genetic correlations among different components of phenotypes, especially resulting from pleiotropy, can constrain or facilitate trait evolution. These factors could especially influence the evolution of traits that are functionally integrated, such as those comprising the flower. Indeed, pleiotropy is proposed as a main driver of repeated convergent trait transitions, including the evolution of phenotypically-similar pollinator syndromes.
- We assessed the role of pleiotropy in the differentiation of floral and other reproductive traits between two species *—Jaltomata sinuosa* and *J. umbellata* (Solanaceae)—that have divergent suites of floral traits consistent with bee- and hummingbird-pollination, respectively. To do so, we generated a hybrid population and examined the genetic architecture (trait segregation and QTL distribution) underlying 25 floral and fertility traits.
- We found that most floral traits had a relatively simple genetic basis (few, predominantly additive, QTL of moderate to large effect), as well as little evidence of antagonistic pleiotropy (few trait correlations and QTL co-localization, particularly between traits of different classes). However, we did detect a potential case of adaptive pleiotropy among floral size and nectar traits.
- These mechanisms may have facilitated the rapid floral trait evolution observed within *Jaltomata*, and may be a common component of rapid phenotypic change more broadly.

## Introduction

One feature of phenotypic evolution is the broad variation in observed rates of trait change, both among traits within a single lineage and among lineages. How quickly phenotypes evolve and in what direction, depends not only on their genetic basis--both the number and effect size of causal loci--and the intensity and nature of selection acting upon them, but also on their associations with other traits. Because of genetic covariance, developmental and phylogenetic constraints, and/or correlated selection pressures (Agrawal & Stinchombe, 2009), phenotypic traits often do not evolve independently from one another. Among these causal mechanisms, strong genetic covariance arises either because different traits have a shared genetic basis (pleiotropy) or because they are based on genes that are physically adjacent and therefore often co-inherited (via linkage). Strong pleiotropy is often proposed to constrain phenotypic evolution by preventing correlated traits from moving efficiently towards their own (different) fitness optima (antagonistic pleiotropy). However, such shared genetic control may also promote phenotypic change if covariation is aligned in the direction of selection (adaptive pleiotropy) (Agrawal & Stinchombe, 2009; Wagner & Zhang, 2011; Smith, 2016). Nonetheless, despite some detailed studies (e.g. Ettensohn, 2013; Shrestha *et al.*, 2014; Manceau *et al.*, 2011; Xu & Schluter, 2015), the genetic architecture of many ecologically important traits remains unclear, including the prevalence of strong genetic associations that could shape the course of phenotypic evolution.

How shared architecture influences phenotypic change is especially relevant for suites of traits that are functionally integrated (Armbruster *et al.*, 2014) -- such as the angiosperm flower (Armbruster *et al.*, 2009) -- because the magnitude and direction of pleiotropy will directly shape if and how these co-varying traits respond to selection. Further, because pleiotropy can constrain or favor particular developmental trajectories, it is often proposed to be a main driver of convergent transitions of integrated traits in different lineages (Preston *et al*., 2011; Smith, 2016). The flower is an especially promising model for assessing the role of pleiotropy in shaping phenotypic evolution. Because flowers mediate fitness through their critical reproductive role, their constituent traits (i.e. reproductive structures, perianth, and other attraction/reward features) are often highly functionally integrated (Conner, 2002; Armbruster *et al.*, 2009). Moreover, repeated transitions of phenotypically similar or convergent suites of floral traits have been identified both within and across groups (Fenster *et al.*, 2004; Goodwillie *et al.*, 2010; Wessinger *et al.*, 2016). For example, multiple parallel shifts from bee-pollination to hummingbird-pollination are associated with parallel transitions to flowers with red petals, large amounts of dilute nectar, and narrow corolla tubes within *Penstemon* (Wessinger *et al.*, 2016); similarly, the evolution of the ‘selfing syndrome’ (i.e. reduced overall size, herkogamy, and floral rewards) often accompanies transitions from outcrossing to predominantly selfing mating systems, and has been documented in multiple lineages (Stebbins, 1974; Goodwillie *et al.*, 2010). Such patterns provide an opportunity to assess the relative frequency of adaptive vs. antagonistic pleiotropy in shaping these repeated trait combinations, within a comparative phylogenetic context.

In addition to these ecological and evolutionary features, the known genetic and molecular bases of floral development (Rijpkema *et al.*, 2006; Smaczniak *et al.*, 2012) themselves suggest that pleiotropy might be a key component shaping floral phenotypic change, as well as provide a functionally-informed framework for identifying how changes in these mechanisms can contribute to the diversification of floral traits. Under the ABC(DE) model of floral development, the combinatorial action of different gene products -- primarily different MADS-box transcription factors -- control transitions to flowering and the specification of floral organ identities and organ maturation, by promoting or repressing different downstream targets (reviewed in O’Maoileidigh *et al.*, 2014; Bartlett, 2017). This combinatorial function, and the ability to regulate shared or partially shared downstream targets, provides a potential mechanistic explanation for strong pleiotropy among floral traits. Moreover, several key regulators of floral development also function during fruit and seed production (Smaczniak *et al.*, 2012; O’Maoileidigh *et al.*, 2014); such correlated effects on fertility traits are another potential way that pleiotropy could shape floral phenotypic evolution.

Several empirical approaches have been taken to assess the genetic architecture underlying floral trait specification and evolution, and to evaluate the strength and direction of pleiotropy. Classical quantitative genetic analyses have revealed varying degrees of genetic covariance among floral traits (Gottlieb, 1984; Conner *et al.*, 2014), while numerous QTL (quantitative trait locus) mapping studies have found that loci for at least some different floral traits appear to co-localize to the same genomic region(s) (reviewed in Smith, 2016). Interestingly, such studies have identified more putative cases of adaptive pleiotropy (e.g. QTL affect more than one trait in the direction of parental trait values) than antagonistic, suggesting that adaptive pleiotropy may be a common mechanism contributing to rapid floral evolution. Because QTL generally span a genomic region that contains more than one gene, such QTL co-localization is consistent with, but not definitive evidence of, a role of pleiotropy in shaping floral trait co-variation (e.g. see Hermann *et al.*, 2013). Nonetheless, identifying strong trait correlations and/or QTL co-localization represents a critical step in assessing how pervasive pleiotropy could be in shaping floral trait evolution.

In this study, we examined phenotypic (co)variation among, and the genetic architecture underlying, floral and other reproductive traits within a segregating hybrid population derived from two *Jaltomata* (Solanaceae) species with divergent floral traits (**Figure 1**). Despite only having diversified with the last 5 million years (Sarkinen *et al.*, 2013; Wu *et al.*, 2018), species of *Jaltomata* are highly phenotypically diverse, including extensive variation in the size, shape, and color of floral traits that is absent among their close relatives within *Solanum* and *Capsicum.* Indeed, phylogenetic analyses suggest numerous transitions in floral traits within the genus, including several instances of convergent evolution (Miller *et al.*, 2011; Wu *et al.*, 2018). Importantly, many of these transitions appear to involve parallel changes in several different traits within a lineage (e.g. from flat corollas with small amounts of lightly colored nectar to highly fused corollas with large amounts of darkly colored nectar), suggesting either a shared genetic basis and/or correlated selection (perhaps pollinator-mediated selection) influences these trait associations.

**Figure 1.**
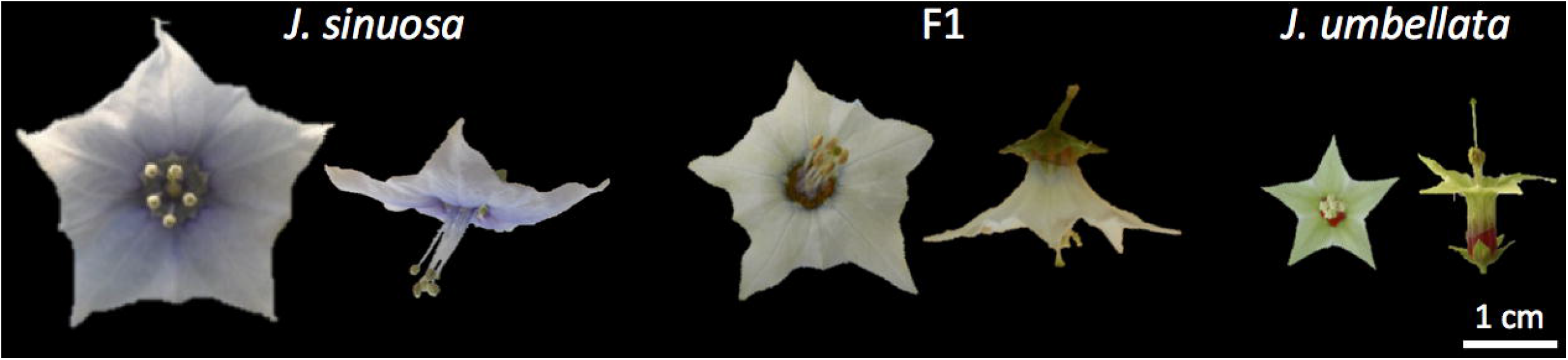
Representative flowers of the parental species and their F1 hybrid.

Here, we identified QTL contributing to floral and other reproductive trait variation within a recombinant population to: 1) examine the genetic architecture underlying reproductive trait divergence; 2) assess the role of genetic linkage and/or pleiotropy (via strong trait correlations and overlapping QTL) in floral trait (co)variation; and 3) assess evidence for a shared genetic basis between different classes of floral trait and other reproductive (specifically fertility) traits. We found evidence for a relatively simple genetic basis underlying most of the examined traits, as well as positive correlations and significant QTL co-localization among traits within each of three trait classes (floral morphology, floral color, and fertility). Together, these features might facilitate rapid changes in these traits. In comparison, we found few associations between traits from different classes, and therefore little evidence for antagonistic pleiotropy among these classes, which otherwise may have constrained trajectories of floral change. One striking exception was an association between flower size and nectar traits that acts in the direction exhibited by multiple species in the genus, suggesting that this could instead be an instance of adaptive pleiotropy.

## MATERIALS AND METHODS

### Study system

*Jaltomata* (Schlechtendal; Solanaceae) is the sister genus to the large and economically important *Solanum* (Olmstead *et al.*, 2008; Sarkinen *et al.*, 2013; Wu *et al.*, 2019), and includes approximately 60-80 species distributed from the Southwestern United States to the Andean region of South America, in addition to several species endemic to the Greater Antilles and the Galapagos Islands (Miller *et al.*, 2011; Mione *et al.*, 2015). Species are highly phenotypically diverse and live in a variety of habitats (e.g. tropical rainforests, rocky foothills, and *lomas* formations).

Here, we focused on a closely related species pair, *J. sinuosa* and *J. umbellata,* that differ in a suite of floral traits that is representative of major floral suites found in other species throughout the genus (**Figure 1; Table S1**). *Jaltomata sinuosa* has large rotate flowers with purple petals and a small amount of concentrated nectar (consistent with bee pollination, Fenster *et al.*, 2004; Mione *et al.*, 2017), while *J. umbellata* has small short-tubular flowers with yellowish-white petals and a large amount of dilute, but dark red, nectar that is visible through the corolla tube (Kostyun & Moyle, 2017). The tubular corollas, large volume of nectar, exerted reproductive organs, and red coloration observed in this species is consistent with hummingbird pollination (Fenster *et al.*, 2004). *Jaltomata sinuosa* is distributed along the Andes in South America from Venezuela to Bolivia, while *J. umbellata* is restricted to lomas formations along the Peruvian coast. Both species are self-compatible (Kostyun & Moyle, 2017) and shrubby, but differ in leaf traits such as overall size and shape, and type of trichomes. This species pair also has several incomplete, intrinsic postzygotic reproductive barriers, including quantitatively reduced fruit set and hybrid seed viability (Kostyun & Moyle, 2017).

### Generation of BC_1_ population and plant cultivation

We developed a segregating hybrid population by crossing *J. sinuosa* and *J. umbellata.* Viable F1 individuals in both directions of the cross produce flowers that are phenotypically intermediate between the parental genotypes (**Figure 1**), and retain reduced but sufficient levels of fertility when back-crossed to parents (Kostyun & Moyle, 2017). Because we were especially interested in the genetic basis of red nectar and a fused corolla tube (exhibited by *J. umbellata,* present but less pronounced in F1s, but absent in *J. sinuosa*; **Figure 1; Figure S7**), we generated the mapping population by backcrossing a single *J. sinuosa* x *J. umbellata* F1 (as the ovule parent) to the original parental *J. sinuosa* individual (as the pollen parent). BC1 individuals were germinated in a growth chamber, and then moved to the Indiana University greenhouse and grown under the same conditions as the parental and F1 individuals (16 hour light cycle, watered twice daily, and fertilized weekly).

### Trait measurements

We measured 25 floral and other reproductive traits within our mapping population, F1, and parental genotypes (**Table 1; Table S2; Figure S1**). Floral morphological traits were measured with hand-held calipers, and nectar volume per flower was measured to the nearest 1 μL using a pipette. To reduce the potential effect of daily environmental variation, nectar volume was always measured in the early afternoon following watering.

**Table 1.** Summary statistics for floral and fertility traits in parental species, F1s, and BC1s. Phenotypic means and variances provided, while significant differences between parental species were assessed with t-tests on transformed data as appropriate (see Methods). Note that differences in seed germination rates were not tested. * p<0.05, ** p<0.001; *** p<0.0001.

| Trait | *J. sinuosa* (n=7) |  | *J. umbellata* (n=5) |  |
| --- | --- | --- | --- | --- |
| Floral Morphology | Mean | Variance | Mean | Variance |
| Buds per Inflorescence | 2.86** | 0.03 | 8.60 | 3.24 |
| Calyx Diameter (mm) | 15.49** | 0.08 | 8.14 | 2.91 |
| Sepal Length (mm) | 7.48** | 0.02 | 4.41 | 0.56 |
| Corolla Diameter (mm) | 29.82*** | 1.92 | 15.32 | 5.70 |
| Corolla Depth (mm) | 1.38** | 0.07 | 10.25 | 5.52 |
| Corolla Fusion (mm) | 9.45 | 0.11 | 10.33 | 1.55 |
| Corolla Fusion Proportion | 0.63* | 0.001 | 0.72 | 0.001 |
| Petal Length (mm) | 14.85 | 0.41 | 14.40 | 0.16 |
| Stamen Length (mm) | 10.80 | 0.29 | 10.45 | 0.22 |
| Anther Length (mm) | 2.03* | 0.02 | 1.73 | 0.04 |
| Ovary Diameter (mm) | 2.28*** | 0.03 | 1.37 | 0.05 |
| Style Length (mm) | 7.90*** | 0.17 | 14.64 | 1.72 |
| Herkogamy (mm) | -2.20*** | 0.01 | 5.37 | 4.62 |
| **Floral Color + Physiology** |  |  |  |  |
| Nectar Color - L | 83.49*** | 20.32 | 50.42*** | 31.63 |
| Nectar Color - a | -1.55*** | 0.95 | 57.61*** | 7.10 |
| Nectar Color - b | 3.47*** | 8.20 | 71.84*** | 59.31 |
| Petal Color - L | 83.18 | 2.59 | 84.15 | 4.86 |
| Petal Color - a | 0.81* | 0.02 | -0.93* | 0.62 |
| Petal Color - b | -1.26*** | 0.02 | 6.71*** | 1.05 |
| Nectar Volume (uL) | 6.86*** | 1.07 | 17.80 | 13.20 |
| **Fertility** |  |  |  |  |
| Viable Pollen Grains | 25000 | 36800000 | 27000 | 60770000 |
| Proportion Viable Pollen | 0.76 | 0.01 | 0.71 | 0.02 |
| Fruit Set | 0.93* | 0.01 | 0.55 | 0.06 |
| Fruit Mass (g) | 0.47** | 0.01 | 0.12 | 0.01 |
| Fruit Diameter (cm) | 1.01* | 0.01 | 0.61 | 0.02 |
| Seed Set | 74.87* | 256.92 | 30.44 | 494.43 |
| Viable Seed Set | 74.87* | 256.92 | 23.59 | 182.31 |
| Viable Seed T50 | 9.33 | -- | 7.00 | -- |
| Viable Seed MGT | 22.00 | -- | 14.00 | -- |

**Table 1 cont.**
| F1s (n=13) |  | BC1s (n=224) |  |
| --- | --- | --- | --- |
| Mean | Variance | Mean | Variance |
| 3.95 | 0.68 | 3.51 | 1.63 |
| 10.55 | 0.48 | 12.10 | 1.08 |
| 5.45 | 0.13 | 6.16 | 0.28 |
| 24.58 | 6.84 | 28.97 | 9.92 |
| 6.64 | 0.58 | 5.22 | 1.35 |
| 9.81 | 0.40 | 9.92 | 1.24 |
| 0.67 | 0.001 | 0.63 | 0.002 |
| 14.65 | 1.61 | 15.78 | 2.88 |
| 10.81 | 0.59 | 10.86 | 1.26 |
| 2.01 | 0.01 | 2.13 | 0.04 |
| 1.62 | 0.02 | 1.82 | 0.04 |
| 12.39 | 1.04 | 10.82 | 1.97 |
| 2.95 | 1.08 | 1.20 | 1.02 |
| 81.31 | 14.35 | 83.99 | 13.42 |
| -0.05 | 101.74 | -4.29 | 14.91 |
| 59.85 | 75.94 | 22.91 | 287.97 |
| 86.93 | 5.82 | 83.56 | 4.60 |
| -0.81 | 0.19 | -0.04 | 0.36 |
| 2.89 | 3.13 | 0.75 | 1.29 |
| 11.90 | 12.32 | 10.76 | 18.87 |
| 22513 | 16602405 | 26886 | 145318716 |
| 0.82 | 0.00 | 0.74 | 0.03 |
| 0.93 | 0.01 | 0.89 | 0.03 |
| 0.24 | 0.01 | 0.29 | 0.01 |
| 0.80 | 0.01 | 0.85 | 0.03 |
| 36.06 | 168.37 | 37.67 | 361.12 |
| 25.46 | 303.95 | 26.28 | 340.08 |
| 34.00 | 1001.78 | 26.70 | 607.37 |
| 46.04 | 622.37 | 39.36 | 458.11 |

Petal and nectar color were quantified using digital photography (Kendal *et al.*, 2013; Garcia *et al.*, 2014): dissected whole petals or nectar drops were photographed on a standard background along with white and black color standards. Light conditions were standardized for all images using RAW Therapee (RAW Therapee Development Team, 2012), and color space attributes were measured in ImageJ (Schneider *et al.*, 2012) as the average value across the entire whole petal or nectar drop sample. Because RGB color attributes are device-dependent (i.e. they can vary depending upon the specific camera used), color values were then converted into device-independent L*a*b color attributes, using the ImageJ Color Space Converter plugin (Schwartzwald, 2012). For petal and nectar color, this produced three color attributes each: Lightness (L; ranges from 0 [black] to 100 [white]), ‘a’ color (more negative values correspond to more green, and more positive values correspond to more magenta), and ‘b’ color (more negative values correspond to cyan, and more positive values correspond to yellow).

We also measured nine fertility traits (**Table 1**) to assess the potential genetic overlap between floral and other reproductive traits, as well as to examine the genetic architecture underlying intrinsic postzygotic barriers between this species pair. Fruit and seed related traits were measured on 2-6 crossed fruit per individual (depending on fruit set). For F1 and BC1 individuals, crossed fruit were produced using pollen from the *J. sinuosa* parental individual. To determine seed germination rates (following Farooq *et al.*, 2005), we soaked 10 seeds per individual in 50% bleach for 30 minutes (to soften the seed coat), rinsed thoroughly, and placed on moist paper within plastic germination boxes. A week after sowing, seed coats were nicked slightly and seeds were given a drop of 10 mM giberellic acid (Sigma) to break dormancy. We then scored germination every 2 weeks for 4 months. Pollen viability was estimated from three different flowers per individual, using established methods (Jewell *et al.*, 2012): Briefly, for each sample all undehisced anthers from a flower were collected into an eppendorf tube containing aniline blue histochemical stain, gently ground with a pestle to release pollen, and viable pollen grains were counted under an EVOS FL Digital Inverted Fluorescence Microscope (Fisher Scientific). From these data, we also calculated the proportion of viable pollen as number of viable pollen grains/total pollen grains in the sample.

### Statistical analyses on phenotypic data

Following Shapiro-Wilk tests to assess normality assumptions, we transformed traits that showed a skewed distribution and/or significantly non-normal residuals. In particular, we arcsine transformed proportional traits (proportion of corolla fusion, fruit set, and proportion of viable pollen), and log-transformed inflorescence size, corolla fusion, petal length, style length, herkogamy, nectar volume, all color attributes, and remaining fertility traits. Significant differences between parental species for all traits were assessed by t-tests (**Table 1**). Distribution plots for all traits are provided in Supplementary Materials (**Figures S3-S6**), while illustrative plots for 15 focal traits are provided in the main text (**Figure 2**). Similarly, phenotypic correlations within the BC1 population were examined among all traits (**Table S3**), while relationships among a subset of focal traits are presented in the main text (**Figure 3**). Given significant correlations among many of the floral morphological traits (**Table S3**), we also used principal component analyses (PCAs) to create separate composite metrics (i.e. Morph PC1-PC3; **Table S4; Figure S2**) as additional measures of floral variation. Trait correlations, including PCs, are provided in **Table S5**, and distribution plots are provided in **Figure S4**.

**Figure 2.**
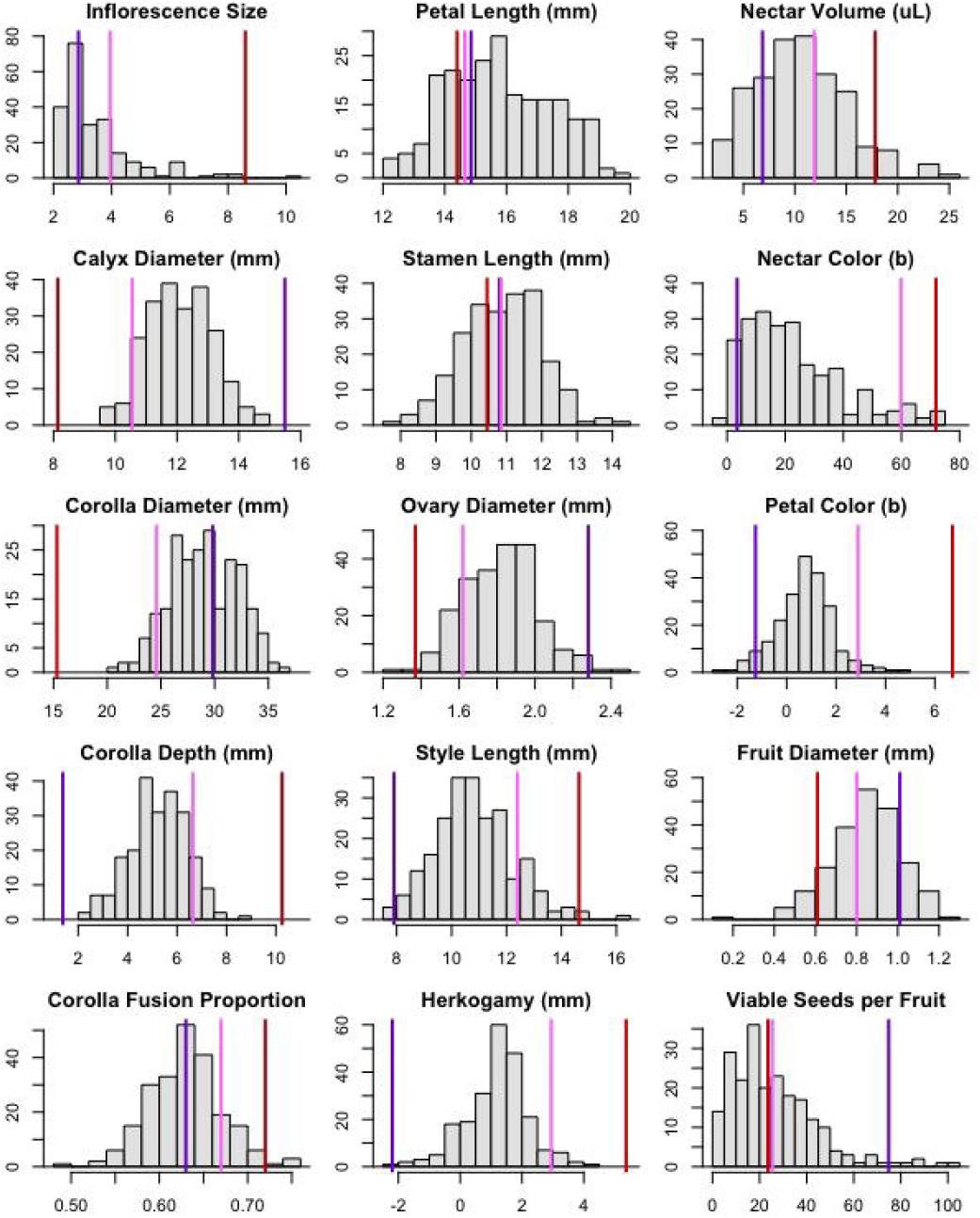
Key trait distributions within the BC1 population, compared to phenotypic means for *J. sinuosa* (purple line), *J. umbellata* (red line), and their F1s (pink line).

### Genotyping and linkage map construction

Genomic DNA was extracted from young leaf tissue from the 2 parental individuals, 13 F1s (including the F1 parent used to generate the population), and 269 BC1s, using the Qiagen DNeasy Plant Mini Kit. DNA quantity and quality were confirmed via Nanodrop (Fisher Scientific) and gel electrophoresis with λ DNA-HindIII Digest marker (New England BioLabs). Samples were then sent to Novogene Corporation (Beijing) for genotyping-by-sequencing (GBS). GBS libraries were prepared using optimized restriction enzymes (MseI and HaeIII), and following insert size selection, sequenced on an Illumina Hi-Seq to generate 150bp paired end reads. Raw reads were trimmed and filtered using Trimmomatic (Bolger *et al.*, 2014), and read quality was checked pre- and post-trimming using fastqc (Andrews, 2010). To identify SNPs, cleaned reads were mapped to the domesticated tomato genome (Tomato Genome Consortium, 2012) using the mem function in BWA (Li, 2013). Alignment files were then input into the Stacks refmap pipeline (Catchen *et al.*, 2013) to determine genotypes. A minimum depth of six identical reads was required to create an individual stack (-m parameter) and automated genotype correction was implemented during export. Subsequent filtering of genotypes was performed during linkage map construction (see below). Reads and genotype data are available in NCBI BioProject PRJNA514267.

To construct the linkage map, we first removed markers that were genotyped in less than 35% of individuals, or showed significant segregation distortion (i.e. alleles at >80% or <20% frequency). The linkage map was constructed using the MST and Kosambi algorithms, implemented in the R package ASMap (Taylor & Butler, 2017). To alleviate map expansion issues, we removed markers which consistently differed from neighboring markers in terms of genotype assignment, indicating a high likelihood of genotyping error. The linkage map was then finalized using the ripple function in R package R/qtl (Broman *et al.*, 2003).

### Identifying QTL

We implemented Haley-Knott regression in R/qtl to identify QTL contributing to each trait. To account for potential environmental contributions to trait variation, we included date of measurement (Month) and location within the greenhouse (Bench) as covariates in our QTL scans. Putative QTL were first identified using the scanone function, followed by permutations for genome-wide LOD significance thresholds. Two dimensional scans (scantwo function) were used in the stepwise qtl function to fit multiple QTL models. These models were used to identify: significant QTL, their 1.5 LOD confidence intervals, their effect sizes (i.e. difference in phenotype mean between homozygotes and heterozygotes), the total amount of phenotypic variance explained by each QTL, the proportion of parental difference explained (relative homozygous effect or RHE), interactions among QTL within traits (testing evidence for epistasis), and potential contributions of covariates. QTL were considered to be co-localized if their 1.5 LOD intervals overlapped. Significant co-localization was assessed by comparing overlap among identified QTL to overlap from 10000 randomly generated distributions, for traits within each category (morphological, color/physiological, or fertility), and between each trait category. Briefly, a custom Python script was used to generate random distributions of QTL (by randomly re-distributing the identified QTL among the 12 linkage groups), and the observed frequency of co-localization in each was recorded for each randomization to generated count distributions, in R. All code used to generate the linkage map, identify QTL, and assess QTL co-location, is available on GitHub (https://github.com/gibsonMatt/jaltomataQTL).

## RESULTS

### Segregation patterns suggest additive alleles underlie most floral traits

Most traits were significantly different between the two parental species (**Table 1**). F1 means were intermediate for most traits as well, except that petals were generally brighter (more white) than either parent, perhaps because F1s had lower amounts of the different pigments found in each parental species (i.e. purple pigment found in *J. sinuosa* petals, and yellow and red pigments found in *J. umbellata)* (**Figures S5+S7**). Other than fruit set and seed germination rates, all traits were unimodally distributed within the BC1s; phenotypic values were intermediate between F1s and the recurrent parent *(J. sinuosa)* for many of these traits, consistent with additive effects (**Figure 2; Figures S3-S6**). Several traits (7 of 25) showed transgressive segregation within the BC1s, including some floral morphological traits, nectar volume and color (all 3 attributes), and seed viability and germination rates (**Table S2; Figures S3-S6**).

### Significant correlations observed within – but generally not between – floral morphology, floral color, and fertility trait categories

Within the BC1s, most traits were not strongly associated with one another. Nonetheless, several correlations remained significant following multiple testing (Bonferroni) correction (**Table S3**), primarily associations that are expected biologically, including allometric relationships among floral organs and positive associations among related fertility traits. For instance, corolla diameter was significantly positively associated with most other morphological traits, suggesting shared genetic control of overall floral size (**Figure 3**), while corolla diameter was also significantly negatively correlated with proportion of corolla fusion (i.e. shorter corolla tubes had wider limbs and longer tubes had narrower limbs, r = −0.348, *p* = 8.68 x 10^-8^). These relationships were recovered with PCA on morphology traits, in which PC1-PC3 explained 76% of the variance among BC1s (**Table S4**). Based on trait loadings, PC1 corresponds to floral width vs. depth, PC2 to overall floral size, and PC3 to relative reproductive organ dimensions. Similarly, related fertility traits also remained strongly correlated, such as fruit mass with seed set, and number of viable pollen grains with proportion of viable pollen (**Table S3**).

**Figure 3.**
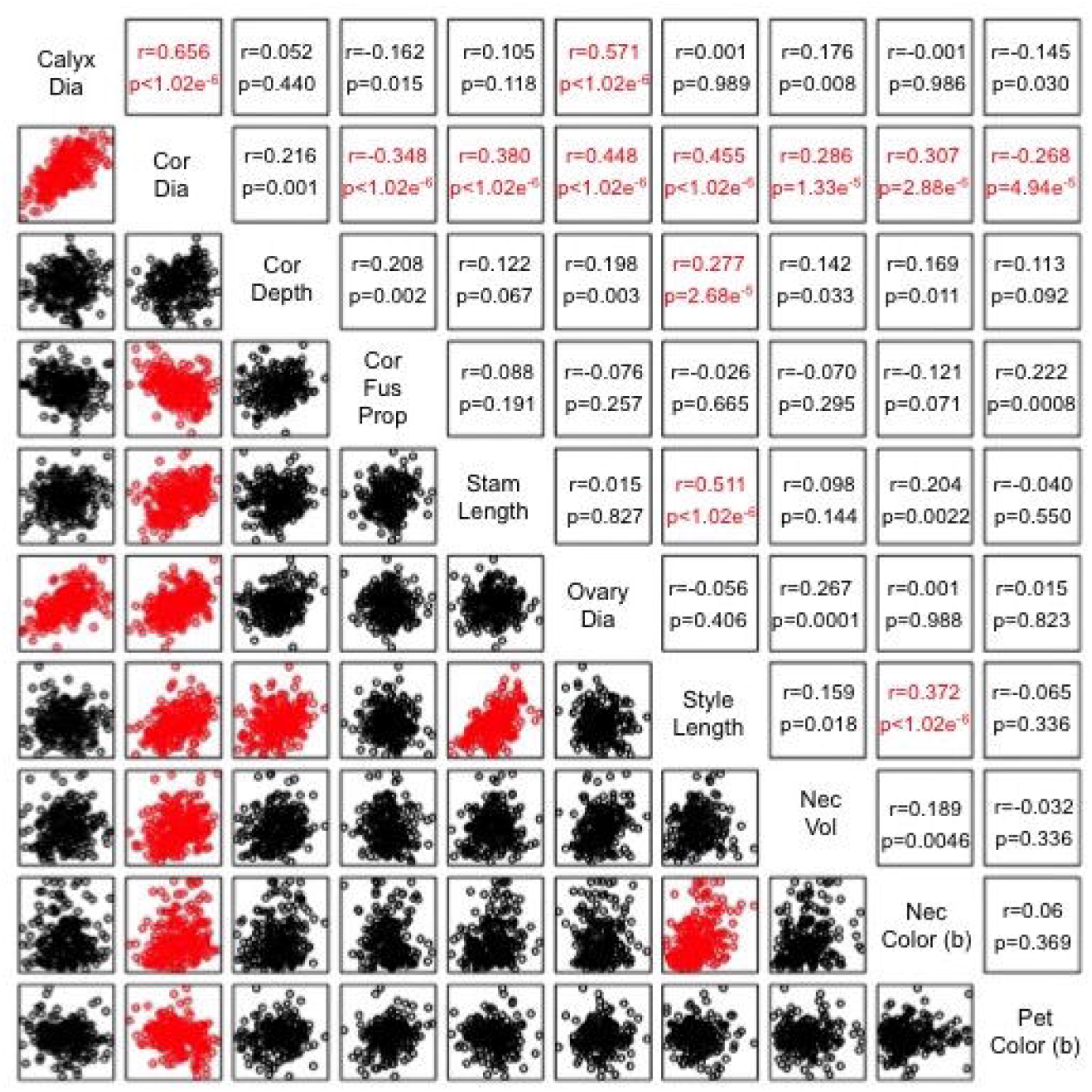
Key floral trait correlations within the BC1 mapping population. Scatterplots are provided below the diagonal, while Spearman’s correlation coefficients and associated p-values are above the diagonal. Statistically significant correlations following Bonferroni correction (p < 5.945 x 10^-5^) are highlighted in red. Correlation values for all traits are provided in Table S3, and for PC traits provided in Table S5.

In contrast, there were relatively few significant correlations among different trait categories. Notable exceptions, however, included a positive relationship between corolla diameter and nectar volume, as well as several morphological traits and certain aspects of color (nectar ‘b’, petal ‘a’ and ‘b’) (**Figure 3; Table S3**). In particular, larger flowers tended to have more red nectar as well as more purple petals. There were also significant positive correlations between pollen viability and each of several components of flower size (as well as Morph PC2 or “size”) (**Tables S3+S5**). This latter relationship seems to be explained by anther size: across 15 *Jaltomata* species, mean viable pollen count is significantly associated with anther size prior to dehiscence (F = 15.56, *p* = 0.0017) (J.L. Kostyun, unpub.).

### Linkage map construction recovered 12 linkage groups

Mapping high quality reads to the tomato genome identified 25,136 SNPs that differentiated the two parental species. Following all subsequent filtering (removing markers genotyped in less than 35% of individuals, with high segregation distortion or non-Mendelian inheritance, or with high likelihood of genotyping errors), we retained 520 high quality markers. Linkage map construction recovered 12 linkage groups (LGs), which correspond to the number of chromosomes in the parental species (Mione *et al.*, 1993; Chiarini *et al.*, 2017). Total map length was 1593.71 cM (65.17 cM – 324.48 cM per chromosome/LG), with an average of 2.92 cM between markers (**Figure 4**).

**Figure 4.**
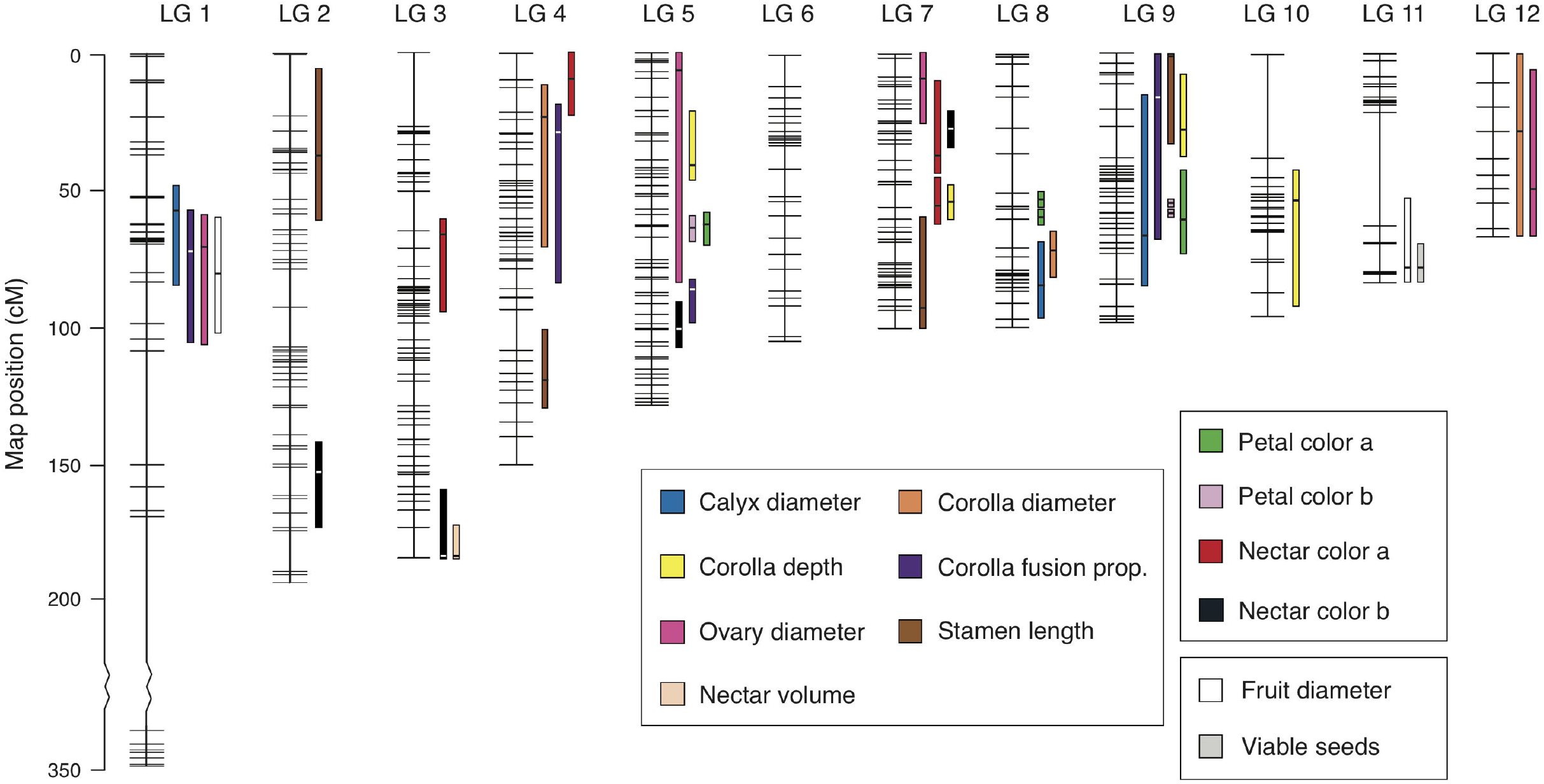
Linkage map and distribution of identified QTL (including 1.5 LOD intervals) for 12 key floral and fertility traits.

### Few moderate to large-effect QTL underlie most traits, with little QTL co-localization between trait categories

We identified QTL for 20 of our 25 examined traits, for a total of 63 QTL (with 4 additional loci for Morph PCs). For traits with identified QTL, most had 2-4 QTL (range=1-7). Alleles at 55 of 67 QTL (82%) acted in a direction consistent with parental values (i.e. the allele from paternal donor *J. umbellata* moved the phenotype of BC1s closer to this species mean); at least one QTL with opposite effects was detected for 10 of the 20 traits (and Morph PC1) (**Table 2; Table S6**). Consistent with trait segregation patterns observed in the BC1, significant interactions (epistasis) among QTL were identified for only two traits: ovary diameter and nectar color ‘a’ (**Table 2; Table S6**).

**Table 2.** QTL for key floral and fertility traits. Includes peak location and LOD of each QTL, 1.5 LOD intervals, proportion phenotypic variance explained, phenotypic effect size and standard error, proportion of parental trait difference explained, and whether the effect is aligned with parental trait values. Comprehensive QTL data, including the full models for all traits, are provided in Table S6. Note: effect sizes are given as the difference between homozygote and heterozygote mean phenotypes (i.e. the effect of a single paternal allele at that QTL), while the proportion of parental trait difference corresponds to the relative homozygous effect (RHE).

| **Trait** | **Linkage Group** | **Peak Location**<br>**(1.5 LOD Interval)** | **Peak LOD** |
| --- | --- | --- | --- |
| *Flower Morphology* |  |  |  |
| Calyx Diameter (3) | 1 | 56 (49-81) | 9.06 |
|  | 8 | 84.3 (69-92.55) | 3.15 |
|  | 9 | 66 (17-85) | 5.59 |
| Corolla Diameter (3) | 4 | 21.1 (13-70) | 3.50 |
|  | 8 | 73.1 (67-84) | 5.87 |
|  | 12 | 37.4 (0-65.17) | 1.57 |
| Corolla Depth (4) | 5 | 40.8 (20-46) | 10.19 |
|  | 7 | 52 (48-60.67) | 17.95 |
|  | 9 | 30 (11-38) | 12.38 |
|  | 10 | 55.8 (41-91) | 4.48 |
| Corolla Fusion Prop. (4) | 1 | 72 (57-106) | 4.60 |
|  | 4 | 26 (19-84) | 3.77 |
|  | 5 | 86 (80.21-99) | 3.57 |
|  | 9 | 16 (0-67) | 7.19 |
| Petal Length (3) | 7 | 100 (25-100.67) | 2.46 |
|  | 8 | 73 (39-92.55) | 3.55 |
|  | 12 | 48.2 (0-65.17) | 2.04 |
| Stamen Length (4) | 2 | 37.5 (6-61) | 4.54 |
|  | 4 | 120.8 (100-133) | 3.77 |
|  | 7 | 91 (59-100.67) | 4.45 |
|  | 9 | 0 (0-33) | 5.32 |
| Anther Length (1) | 1 | 18 (8-35.87) | 3.78 |

**Table 2. cont.**
|  |  |  |  |
| --- | --- | --- | --- |
| Ovary Diameter (4) | 1 | 68 (60-107) | 5.77 |
|  | 5 | 72 (0-84) | 2.53 |
|  | 7 | 9.9 (0-25) | 3.56 |
|  | 12 | 48.2 (5-65.17) | 4.31 |
| Style Length (2) | 2 | 95.2 (55-126) | 2.79 |
|  | 9 | 22 (1-30) | 9.89 |
| *Flower Physiology* |  |  |  |
| Nectar Volume (1) | 3 | 187.2 (172-187.19) | 2.81 |
| *Flower Color* |  |  |  |
| Nectar L (2) | 2 | 154.9 (147-171.87) | 2.97 |
|  | 5 | 103.9 (74-115.35) | 6.44 |
| Nectar a (4) | 3 | 66.1 (60-95) | 7.69 |
|  | 4 | 14 (0-23.74) | 3.99 |
|  | 7 | 29.1 (10-42) | 2.32 |
|  | 7 | 56.2 (43-63) | 3.54 |
| Nectar b (4) | 2 | 152 (141-171.87) | 4.50 |
|  | 3 | 186 (164-187.19) | 2.83 |
|  | 5 | 100 (94-111) | 11.00 |
|  | 7 | 26 (20.02-32) | 9.02 |
| Petal L (2) | 4 | 23.7 (0-81) | 2.40 |
|  | 5 | 65.3 (58-71) | 3.83 |
| Petal a (4) | 5 | 62 (58-68) | 12.11 |
|  | 8 | 52 (49-52.48) | 8.54 |
|  | 8 | 52.5 (52-53) | 8.13 |
|  | 9 | 64.1 (44-76) | 9.89 |
| Petal b (3) | 5 | 64 (60-69) | 21.46 |
|  | 9 | 52 (52-54) | 5.60 |
|  | 9 | 55.8 (54.05-57) | 4.22 |
| *Fertility* |  |  |  |
| Fruit Diameter (2) | 1 | 80 (61-103) | 4.67 |
|  | 11 | 76.9 (51-79.93) | 3.09 |
| Fruit Mass (4) | 1 | 96 (60-146) | 5.95 |
|  | 3 | 170 (48-187.19) | 2.99 |
|  | 11 | 76.5 (54-79.93) | 4.19 |
|  | 12 | 0 (0-10.64) | 4.66 |
| Seed Set (2) | 11 | 76 (63-79.93) | 6.64 |
|  | 12 | 0 (0-63) | 2.86 |
| Viable Seed Set (1) | 11 | 76.9 (68-79.93) | 4.42 |

**Table 2. cont.**
| **%PVE** | **Effect ± SE** | **%Parental**<br>**Trait**<br>**Difference**<br>**(RHE)** | **Aligned**<br>**with**<br>**Parental**<br>**Value?** |
| --- | --- | --- | --- |
| 13.67 | -0.803 ± 0.120 | 0.22 | Y |
| 4.46 | -0.457 ± 0.120 | 0.12 | Y |
| 8.13 | -0.598 ± 0.116 | 0.16 | Y |
| 4.47 | 1.387 ± 0.344 | -0.19 | N |
| 7.69 | -1.902 ± 0.360 | 0.26 | Y |
| 1.96 | -0.963 ± 0.361 | 0.13 | Y |
| 11.20 | 0.844 ± 0.119 | 0.19 | Y |
| 21.47 | 1.146 ± 0.116 | 0.26 | Y |
| 13.94 | 0.942 ± 0.119 | 0.21 | Y |
| 4.64 | -0.521 ± 0.114 | -0.12 | N |
| 7.02 | -0.022 ± 0.005 | -0.56 | N |
| 5.70 | -0.021 ± 0.005 | -0.53 | N |
| 5.39 | 0.022 ± 0.005 | 0.56 | Y |
| 11.27 | 0.029 ± 0.005 | 0.74 | Y |
| 3.29 | 0.018 ± 0.005 | -2.66 | N |
| 4.78 | -0.022 ± 0.005 | 3.32 | Y |
| 2.71 | -0.017 ± 0.005 | 2.50 | Y |
| 5.80 | -0.548 ± 0.119 | 3.18 | Y |
| 4.78 | -0.499 ± 0.119 | 2.90 | Y |
| 5.68 | 0.551 ± 0.121 | -3.20 | N |
| 6.86 | 0.619 ± 0.124 | -3.60 | N |
| 6.48 | -0.116 ± 0.027 | 0.78 | Y |

**Table 2. cont.**
|  |  |  |  |
| --- | --- | --- | --- |
| 9.21 | $-0.120 \pm 0.023$ | 0.26 | Y |
| 3.90 | $0.092 \pm 0.027$ | -0.20 | N |
| 5.56 | $-0.094 \pm 0.023$ | 0.21 | Y |
| 6.78 | $-0.104 \pm 0.023$ | 0.23 | Y |
| 3.80 | $0.022 \pm 0.006$ | 0.16 | Y |
| 14.51 | $0.044 \pm 0.006$ | 0.33 | Y |
| 5.22 | $0.098 \pm 0.027$ | 0.48 | Y |
| 5.12 | $-0.008 \pm 0.002$ | 0.08 | Y |
| 11.51 | $-0.014 \pm 0.003$ | 0.13 | Y |
| 13.04 | $0.062 \pm 0.010$ | 0.23 | Y |
| 6.51 | $-0.043 \pm 0.010$ | -0.16 | N |
| 3.72 | $-0.040 \pm 0.012$ | -0.15 | N |
| 5.74 | $0.050 \pm 0.012$ | 0.18 | Y |
| 5.25 | $0.161 \pm 0.036$ | 0.27 | Y |
| 3.24 | $0.138 \pm 0.038$ | 0.23 | Y |
| 13.74 | $0.288 \pm 0.039$ | 0.48 | Y |
| 11.03 | $0.230 \pm 0.035$ | 0.39 | Y |
| 4.43 | $0.005 \pm 0.002$ | 2.03 | Y |
| 7.19 | $0.007 \pm 0.002$ | 2.92 | Y |
| 15.28 | $-0.122 \pm 0.016$ | 0.44 | Y |
| 10.35 | $-1.355 \pm 0.210$ | 4.81 | Y |
| 9.82 | $1.312 \pm 0.209$ | -4.65 | N |
| 12.18 | $-0.097 \pm 0.014$ | 0.35 | Y |
| 32.83 | $0.210 \pm 0.019$ | 0.56 | Y |
| 7.20 | $0.486 \pm 0.094$ | 1.31 | Y |
| 5.35 | $-0.415 \pm 0.093$ | -1.11 | N |
| 8.64 | $-0.026 \pm 0.005$ | 0.52 | Y |
| 5.61 | $-0.020 \pm 0.005$ | 0.40 | Y |
| 9.35 | $-0.026 \pm 0.005$ | 0.44 | Y |
| 4.56 | $-0.018 \pm 0.002$ | 0.31 | Y |
| 6.46 | $-0.021 \pm 0.005$ | 0.35 | Y |
| 7.22 | $-0.023 \pm 0.005$ | 0.40 | Y |
| 12.28 | $-0.157 \pm 0.028$ | 0.65 | Y |
| 5.07 | $-0.109 \pm 0.030$ | 0.45 | Y |
| 8.85 | $-0.211 \pm 0.046$ | 0.75 | Y |

The amount of phenotypic variation explained by each QTL ranged from 2-33% (**Table 2; Table S6**). Of these, 19 traits (and Morph PC1 and PC3) had at least 1 moderate effect QTL (defined as explaining 5-25% of the variance), and 1 trait (petal color ‘b’) had a QTL with major effect (defined as >25% variance explained) (**Table 2; Table S6**). The lower bound of this range suggests that we had reasonable power to identify QTL with even relatively small effects -- explaining as little as 2% of the variance. We note however, that like most QTL analyses (Broman, 2001), we likely have limited power to detect all QTL contributing to our examined traits. Nonetheless, under strict additivity, QTL account for >50% of the parental difference (relative homozygous effect, RHE) in 14 of the 20 traits for which we identified QTL (as well as Morph PC1-3); for 8 of these traits (as well as Morph PC1-3), >75% of the parental difference is accounted for (**Table 2; Table S6**). That is, for most of our traits, detected loci account for the majority of parental phenotypic differences. In contrast, for four traits (corolla diameter, proportion of corolla fusion, herkogamy, and inflorescence size) QTL accounted for <20% of the parental difference, suggesting additional undetected loci contribute to these traits. Finally, we estimated RHE >1 for 5 floral traits that also show the most transgressive segregation among BC1s (nectar color ‘b’, petal color ‘L’, petal length, corolla fusion, and Morph PC2) (**Table S6; Figures S3-5**).

Although every linkage group had at least one QTL, these were not distributed uniformly across the genome, with notable clusters on LG1 and LG9 (**Figure 4; Table 2; Tables S6+S8**). We also identified several instances of QTL co-localization within trait categories, especially for morphology and fertility traits, which each had significantly more cases of co-localization (1.5 LOD overlap) than expected by chance (133 observed vs. upper bound of 115 expected overlaps, *p* = 6.9 x 10^-4^; 8 observed vs. upper bound of 4 expected overlaps, *p* = 5.0 x 10^-6^, respectively) (**Table S7; Figure S8**). These instances of co-localization included QTL for biologically-related traits for which we observed strong correlations, such as those underlying petal length and corolla diameter on LG8 and LG12, corolla depth and corolla fusion on LG7, LG9 and LG10, and fruit mass and seed set on LG11 and LG12 (**Tables S3+S8**). In some cases, co-localized QTL share the same or a very close peak marker (e.g. calyx diameter and inflorescence size on LG1, and petal length, corolla fusion, and ovary diameter on LG12; **Tables S6+S8**), which suggests pleiotropic effects, however we note that the large 1.5 LOD intervals of some QTL will increase instances of incidental co-localization events.

In contrast, co-localization between different trait categories was never greater than expected by chance (**Table S7; Figure S8**). This is consistent with mostly incidental overlap between QTL for traits in different categories. Nonetheless, we did detect co-localization at the same or very close peak markers for three sets of biologically interesting trait combinations: corolla diameter, proportion of corolla fusion, and petal color ‘L’ on LG4; corolla fusion, corolla depth, and nectar color ‘a’ on LG7; and nectar volume and nectar color ‘b’ on LG3 (**Figure 4; Tables S6+S8**). Not all of the traits within each of these sets is significantly phenotypically correlated in the BC1 (**Table S3**), especially correlations involving nectar color ‘a’ and ‘b’, possibly because these nectar color traits have additional QTL that are not shared with the corolla traits or nectar volume. Nonetheless, the colocalized QTL still explain large fractions of the parental difference for these traits: on LG7 the QTL colocalized with corolla fusion and depth explains ~20% of the parental difference in nectar color ‘a’, while the potentially shared QTL on LG3 explains ~25% of the parental difference in nectar color ‘b’ and nearly 50% of the parental difference in nectar volume (**Table 2; Table S6**). Together, these provide intriguing cases of potential adaptive pleiotropy, such that alleles at QTL may have simultaneously acted to increase corolla tube depth and nectar darkness, and increase nectar production and nectar darkness, respectively.

## DISCUSSION

Genetic correlations among different components of the phenotype, especially resulting from pleiotropy, can constrain or facilitate trait evolution (Agrawal & Stinchcombe, 2009). Pleiotropy could have particularly strong effects on the evolution of traits that are functionally integrated, such as those comprising the flower (Armbruster *et al.*, 2009; Smith, 2016). Here, we examined the genetic architecture underlying floral trait evolution within florally diverse *Jaltomata,* including whether pleiotropy might have shaped observed variation. We found that most of our examined traits have a relatively simple genetic basis, with few to moderate QTL with largely additive effects. We also identified strong correlations and significant QTL overlap within trait categories, but few associations across different types of traits. The exceptions however, between certain aspects of floral morphology and nectar traits (volume, and both ‘a’ and ‘b’ color), are consistent with existing trait associations that are observed across the genus, suggesting that these could be examples of adaptive pleiotropy. Although we did not directly assess the role of selection, our data indicate that the rapid floral trait evolution observed in this group could have been facilitated by a relatively simple genetic basis for individual floral traits, and a general absence of antagonistic pleiotropy among different types of reproductive traits.

### Few genetic changes could underlie floral trait shifts

The relatively simple genetic architecture that we detect for most of our floral traits might be one mechanism that has permitted rapid floral evolution within the genus. Indeed, our inference that few QTL underlie corolla depth agrees with comparative development data from these species (Kostyun *et al.*, 2017) in which we observe that relatively simple heterochronic changes in corolla trait growth rates distinguish these rotate vs. tubular corolla forms. Interestingly, our findings are also consistent with previous studies of floral trait genetics between closely related species (Smith, 2016), as well as a quantitative genetics study of floral trait variation between two ecotypes of *Jaltomata procumbens* (Mione & Anderson, 2017). In particular, Mione and Anderson (2017) estimated 1-4 loci contributed to various floral traits, as well as petal spot patterning. In other systems, one or few QTL have been found for species differences in nectar volume (e.g. Bradshaw *et al.*, 1998; Stuurman *et al.*, 2004; Wessinger *et al.*, 2014; but see Nakazato *et al.*, 2013), similar to our inference of a single QTL for this trait. For petal and nectar color (including Lightness, color ‘a’ and color ‘b’), we identified 4 and 7 QTL, respectively, with at least 1 QTL per trait with moderate or large effects (i.e. 10 - 33% PVE) (**Table 2; Table S6**), similar to other systems that generally identify few loci of large effect for petal color (e.g. Bradshaw *et al.*, 1998; Wessinger *et al.*, 2014).

Perhaps unlike these cases, however, it is likely that loci controlling color differences in *Jaltomata* are regulators of pigment quantity rather than presence/absence biosynthesis, because both nectar and petal color show gradation in the BC1s rather than discrete color bins. Based on synteny with domestic tomato, our identified petal color QTL occur on the same chromosomes as several known anthocyanin biosynthesis genes (i.e. DFR on ch2, CHS1 and F3H on ch9, and ANS1 on ch10, in tomato), but do not appear to overlap with the chromosomal locations of these genes. Preliminary data from a VIGS (virus-induced gene silencing) pilot study in *J. sinuosa* do indicate that the purple petal pigment is an anthocyanin however (J.L. Kostyun and J.C. Preston, unpub.), whereas for nectar color, preliminary data suggest that an indole-flavin contributes to red pigment in *J. umbellata* (J.L. Kostyun and D. Haak, unpub.), consistent with our inference that color variation is unassociated between these different floral components.

In addition to relatively few contributing loci, many of the examined floral traits also appear to be underpinned by additive effects (**Table 1; Figure 2; Figures S3-S6**), while epistatic effects were comparatively rare (see below). Both are factors that might also facilitate more rapid responses to selection. Other studies have similarly found that floral size traits are often additive (Gottlieb, 1984), including within *Jaltomata* (Mione & Anderson, 2017). Although several floral traits showed transgressive segregation within our BC1s, which could indicate epistatic interactions, similar patterns can result from unique combinations of additive alleles that have opposite effects in the parental species (e.g. deVicente & Tanksley, 1993) and we identified individual QTL with these opposing effects for all traits with transgressive segregation (**Table S6; Figures S3-5**). In addition, our models only detected significant interactions among QTL for ovary diameter and nectar color ‘a’ (**Table S6**), consistent with a general lack of epistatic interactions for floral traits. Therefore, while we may not have uncovered all epistatic interactions contributing to our traits--because some QTL might be undetected in our study, and our models did not assess between trait interactions--both QTL and overall segregation patterns suggest that floral traits are primarily underpinned by additive rather than epistatic effects.

In contrast, many of the fertility traits had segregation patterns consistent with epistatic interactions. BC1 individuals tended to have lower seed set and poorer quality seeds (decreased viability and response to germination-inducing stimuli), and the recombinant BC population contained a subset of highly sterile individuals. The segregation of recombinant individuals with reduced viability and fertility often occurs in hybrids (Baack *et al.*, 2015), including in hybrids from additional *Jaltomata* species pairs (Kostyun & Moyle, 2017). Such patterns are typically due to deleterious epistatic interactions between loci that have diverged between the two parental lineages, as has been shown in close relatives including tomatoes (Moyle & Nakazato, 2008; Sherman *et al.*, 2014). These observations in *Jaltomata* are similarly consistent with a specific role for epistasis among incompatible alleles in the expression of postzygotic reproductive isolation.

### Reduced constraints may also have facilitated rapid floral trait evolution

Because rapid floral evolution may occur either through a lack of antagonistic pleiotropy or through adaptive pleiotropy, we assessed evidence for these potential mechanisms within *Jaltomata*. Within trait categories, we detected positive but modest associations between several floral size traits, and among biologically related fertility traits (e.g. fruit size and seed set) (**Figure 3; Table S3**), as well as significant co-localization of QTL for these groups of traits (**Tables S7-S8**). Morphological associations in particular suggest that shared growth regulators (e.g. Sicard & Lenhard, 2011; Brock *et al.*, 2012) contribute to observed variation in floral organ sizes, but do not completely determine it; that is, overall floral size appears to share growth regulators (potentially represented by co-localized QTL among floral size traits), but different floral organs also appear to have unique growth regulator(s) (consistent with unique QTL). In contrast, we detected fewer instances of strong trait correlations and QTL co-localization between different trait categories (Tables S3+S7-S8). This general lack of antagonistic pleiotropy among different classes of floral and fertility traits may have facilitated rapid floral evolution in this system by minimizing constraints on the available combinations of floral traits.

Despite this general pattern, we did identify several instances of QTL co-localization that might represent adaptive pleiotropy, specifically among different corolla and nectar traits. Overall, these associations would have produced flowers with larger corolla limbs, deeper corolla tubes, and a greater volume of darker colored nectar (**Table 2; Figure 4; Tables S6+S8**). Interestingly, this trait combination (large flowers with copious dark nectar) is actually not exhibited by either parental species used in this experiment (**Figure 1; Table 1**); however, it is found in numerous other *Jaltomata* species and is consistent with suspected pollination syndromes within the genus (see below; Miller *et al.*, 2011; Kostyun & Moyle, 2017). Correlated changes in corolla size and nectar volume have also been observed in several other systems, including those with documented or hypothesized pollinator shifts (e.g. *Petunia, Mimulus,* and *Penstemon;* Galliot et al. 2006; reviewed in Smith, 2016).

### *Ecological context for rapid floral change in* Jaltomata

Overall, our findings suggest potential mechanistic explanations for the evolution of remarkable floral trait diversity among *Jaltomata* species within the last 5 million years (Sarkinen *et al.*, 2013). Traits with a relatively simple genetic basis that are uncoupled from other floral and fertility traits have fewer mechanistic constraints, and therefore can more rapidly respond to selective opportunities as they arise (given sufficient standing or new genetic variation) (Agrawal & Stinchcombe, 2009; Smith, 2016). Although we have not yet directly assessed the role of selection in shaping floral differences among species, several features of *Jaltomata* floral biology are consistent with pollinator-mediated selection on floral traits (van der Niet & Johnson, 2012), likely in conjunction with mating-system related changes (Goodwillie *et al.*, 2010). First, floral trait variation within *Jaltomata* shows clear hallmarks of selection imposed by pollinator differentiation. Of the two main lineages within the genus, nearly all species in the Central and South American clade have relatively small, ancestrally rotate flowers with small amounts of lightly colored nectar (Wu et al. 2018), and hymenopterans have been observed visiting several of these species in their native range (T. Mione, pers. comm.). In contrast, many species in the clade that is restricted to South America—including *J. umbellata* examined here—have floral features associated with attraction of vertebrate pollinators (Fenster *et al.*, 2004). Several of these species with larger flowers, a highly fused corolla (either campanulate or tubular), and copious amounts of darkly colored nectar are visited by hummingbirds (T. Mione, per. comm.); intriguingly, this repeated natural trait covariation is consistent with the genetic association between floral size and nectar traits we identified here.

These features indicate that shifts among different types of pollinators are a likely source of selection for floral differentiation among species within *Jaltomata,* however, mating system variation in the genus may also have facilitated these transitions. Self-incompatibility is the ancestral state in the Solanaceae (Steinbachs & Holsinger, 2002) and is broadly persistent in close relatives *Solanum* and *Capsicum* (Goldberg *et al.*, 2010). In contrast, all examined *Jaltomata* species are self-compatible (Mione, 1992; Kostyun & Moyle, 2017; J.L. Kostyun & T. Mione, unpub.), indicating that gametophytic self-incompatibility was lost early in the evolution of this clade. The presence of delayed selfing and strong herkogamy in many species (e.g. Mione *et al.*, 2015; Mione *et al.*, 2019), in addition to field observations of pollinators (above; T. Mione, pers. comm.), indicate that species most likely employ a mixed mating strategy in their native ranges. Mixed mating strategies are generally observed to maintain the largest amount of floral trait variation, compared to predominant selfing or enforced outcrossing (Goodwillie *et al.*, 2005; Rosas-Guerrero *et al.*, 2014). In addition, they have been predicted to facilitate pollinator shifts--especially to pollinators that might be more efficient but potentially unreliable (such as hummingbirds)—because they allow reproductive assurance (via selfing, when pollinators are limited) and increase the expression of new floral trait variation controlled by recessive alleles (Goodwillie *et al.*, 2005; Brys *et al.*, 2013; Wessinger & Kelly, 2018). Notably, our data indicate that dark/red colored nectar is at least partially recessive (**Figure 2**), and that red petal pigmentation is completely recessive (**Figure S7**), consistent with this novel variation being based on new recessive alleles. Together, these observations are intriguing as they suggest that mixed mating, in conjunction with the specific genetic architecture of floral trait variation, might have facilitated the evolution of new floral trait variation in *Jaltomata*.

## CONCLUSIONS

Genetic correlations among floral traits, especially those due to pleiotropic effects, likely shape permitted trajectories of floral evolution. To assess how such genetic associations might have contributed to observed patterns of floral diversity in *Jaltomata,* we examined segregation patterns and genetic architecture of 25 floral and fertility traits in a hybrid (BC1) population generated from parents with divergent floral traits. Our data are consistent with several mechanisms that could have allowed rapid floral trait evolution in this system: a largely simple genetic basis underlying variation in most of our floral traits, a general absence of antagonistic pleiotropy constraining floral evolution, and a potential instance of adaptive pleiotropy governing floral size and nectar traits. This genetic architecture, in combination with pollinator-mediated selection on a background of self-compatible mixed mating, might have uniquely positioned this genus for the rapid floral diversification now evident within *Jaltomata*.

## ACKNOWLEDGEMENTS

We thank the IU greenhouse staff for plant care, CJ Jewell and David Haak for logistical support, undergraduate research assistants (especially Meret Thomas-Huebner, Devki Shukla, and Shachia Jackson) for assistance with data collection, Tim Leslie for assistance with simulations, and Mark Rausher and two anonymous reviewers for feedback on a previous version of the manuscript. This work was supported by the IU Biology Department, National Science Foundation Award (NSF DEB 1136707) to LCM, and National Science Foundation Graduate Research Fellowship Program (NSF DEB 1342962) and Doctoral Dissertation Improvement Grant (NSF DEB 1601078) to JLK.

## AUTHOR CONTRIBUTIONS

JLK and LCM designed the experiment, JLK generated experimental materials, JLK and CMK collected phenotypic data, JLK and MJSG analyzed the data, and JLK and LCM wrote the paper with input from CMK and MJSG.

## Supplementary Information

**Table S1**. Accession information for material used and generated in this study.

**Table S2**. Trait measurements for all individuals phenotyped.

**Table S3**. Correlations among all measured traits within BC1 individuals.

**Table S4**. Principal component loadings for highly correlated traits.

**Table S5**. Correlations among PC traits within BC1 individuals

**Table S6**. Full QTL models for all examined traits.

**Table S7**. Expected and observed counts of QTL overlap.

**Table S8**. Identified instances of QTL co-localization, based on overlapping 1.5 LOD intervals.

**Figure S1**. Measured morphological traits on mature flowers.

**Figure S2**: Biplots following PCA on floral morphological traits

**Figures S3-S4**: Distributions for floral morphological traits within the mapping population.

**Figure S5**. Distributions for physiological and color traits within the mapping population.

**Figure S6**. Distributions for fertility traits within the mapping population.

**Figure S7**. Representative examples of petal and nectar color variation among F1 individuals, compared to either parent.

**Figure S8**. Comparison of observed number of QTL overlaps with counts from 10000 randomly generated simulations, for QTL co-localization overlap within and between trait categories (morphology, color, and fertility).

